# LSM14B is essential for mitochondrial clustering in the oocyte meiosis

**DOI:** 10.1101/2023.04.24.538190

**Authors:** Yanling Wan, Shuang Yang, Tongtong Li, Yuling Cai, Mingyu Zhang, Tao Huang, Yue Lv, Gang lu, Jingxin Li, Qianqian Sha, Zijiang Chen, Hongbin Liu

## Abstract

As oocyte meiotic maturation, they undergo two successive meiotic M phases, notably lacking an intervening interphase phase. During these M phases, oocytes remain transcriptionally quiescent, and we now know that “translational repressed mRNAs” are stored in a structure called the mitochondria associated ribonucleoprotein domain (MARDO). LSM14B is one of the abundant proteins of MARDO, and is predicted to bind mRNA, but its function(s) remain elusive. Here, we demonstrate that LSM14B functions to promote MARDO assembly in mouse oocytes. We also found that LSM14B knockout female mice are infertile, and show that the knockout oocytes fail to enter meiosis II, instead entering an aberrant interphase-like stage. Finally, we show that the failure of oocyte maturation results from decreased expression of Cyclin B1. Our study has revealed that the RNA-binding protein LSM14B modulates MARDO assembly and is essential for oocyte meiotic maturation.

## Introduction

Mammalian oocytes of high-quality are required for successful reproduction. Mice oocytes arrest at the first meiotic prophase, known as the germinal vesicle (GV) stage. Based on chromatin morphology, the GV oocytes can be categorized as surrounded nucleolus (SN) or nonsurrounded nucleolus (NSN) oocytes. The process of oocyte meiotic maturation begins with meiotic resumption via GV breakdown (GVBD)[1]. After polar body 1 (PB1) extrusion occurs, oocytes enter meiosis II without going through the S phase and arrest at metaphase II (MII) until fertilization[2–4]. The activation of maturation-promoting factor (MPF) is required for entrance to meiosis II, and Cyclin B1 is understood as the regulatory subunit of MPF[5, 6]. After meiotic resumption, the oocyte is transcriptionally quiescent [7]; the oocyte can only use stored mRNAs to synthesize new proteins, with translational activation controlled by post-transcriptional regulation of maternal mRNAs[9].

Approximately 90% of the energy requirements for oocyte maturation are met by production of adenosine triphosphate (ATP) through oxidative phosphorylation (OXPHOS) reactions in mitochondria[10]. At metaphase I (MI), oocyte mitochondria cluster on the periphery of the spindle during the so-called ‘ring stage’, and this clustering is mediated at least in part through adaptive cytoplasmic filamentous actin (F-actin) network activity[10–12]. Subsequently, there is a gradual dispersion of mitochondria as oocytes transition from meiosis I to meiosis II[19]. Mitochondrial clustering defects have been linked to a range of devastating abnormities during oocyte maturation[13, 14], yet knowledge about how mitochondria cluster and then disperse during oocyte maturation remains limited.

The RNA-binding protein ZAR1 was shown to co-localize with mitochondria in oocytes, and ZAR1 (as well as other RNA-binding proteins including DDX6 and YBX2) are present in a structure called a mitochondria associated ribonucleoprotein domain (“MARDO”)[19], which functions as a storage site for “translationally repressed mRNAs” in oocytes of various mammalian species[19]. MARDO is first evident in NSN oocytes, and then grows larger with an increasing mitochondrial membrane potential, becoming most prominent in MI oocytes[19]. Concomitant with the dispersion of mitochondria clustering, MARDO dissolution occurs as oocytes transition from meiosis I to meiosis II; however, it should be stressed that MARDO dissolution occurs independently of cell cycle progression *per se*[19].

LSM14 also named RAP55, this 55kDa RNA-binding protein is evolutionarily conserved across vertebrate species[15], most of which have two paralogous proteins: LSM14A and LSM14B[16, 17]. Studies in HeLa cells showed that LSM14 is present in P-bodies (which function in storing mRNAs)[18]. In mitosis, LSM14 has been reported to regulate the mitotic G2/M transition as RNA-binding translational repressors [16, 19]. A previous study reported that knockdown of LSM14A in oocytes had no effect on oocyte maturation, whereas knockdown of LSM14B resulted in metaphase I (MI) arrest[20]. Although LSM14B is one of the most abundant proteins in MARDO, the physiological function(s) of it remain unknown.

Here, fertility testing showed that female LSM14B knockout mice are infertile, and a subsequent detailed analysis isolated the reproductive defect to the meiosis I and II transition stage of oocyte meiotic maturation. Cell biology analysis revealed that LSM14B knockout oocytes have an abnormal distribution of mitochondria at the GV and MI stage. We show through microinjection experiments that LSM14B functions in the assemble of MARDO in MI oocytes.

Working with LSM14B knockout mice, we demonstrate that LSM14B is specifically expressed in oocytes and is essential for the meiosis I to meiosis II transition in mouse oocytes. By co-immunoprecipitation and microinjection, we provide evidence that LSM14B co-related with ZAR1 in an RNA-mediated manner and promotes MARDO assembly in MI oocytes. Through the integration analysis of transcriptomics and proteomics then validation by immunoblotting, we confirmed that in LSM14B knockout mice, of the expression of *Cyclin B1* was downregulated. This study uncovers previously unstudied roles of LSM14B in promoting MARDO assembly and female reproduction.

## Results

### Profiling LSM14B expression in oocytes and early embryos

LSM14B is highly expressed in oocytes and LSM14B knockout in mice and causes female infertility. We profiled *Lsm14b* mRNA levels in diverse somatic tissues, oocytes, and early embryos by qPCR. The *Lsm14b* mRNA levels were obviously higher in oocytes than in somatic tissues (Figure1A). We noted a trend wherein the *Lsm14b* mRNA level is highest in germinal vesicle (GV) stage oocytes, lower in zygotes, and extremely low or absent in 2-cell embryos (Figure1A). Immunoblotting against LSM14B showed high accumulation in GV oocytes, with the abundance decreasing during meiotic maturation and becoming marginal in 2-cell embryos (Figure 1B).

**Figure 1.**
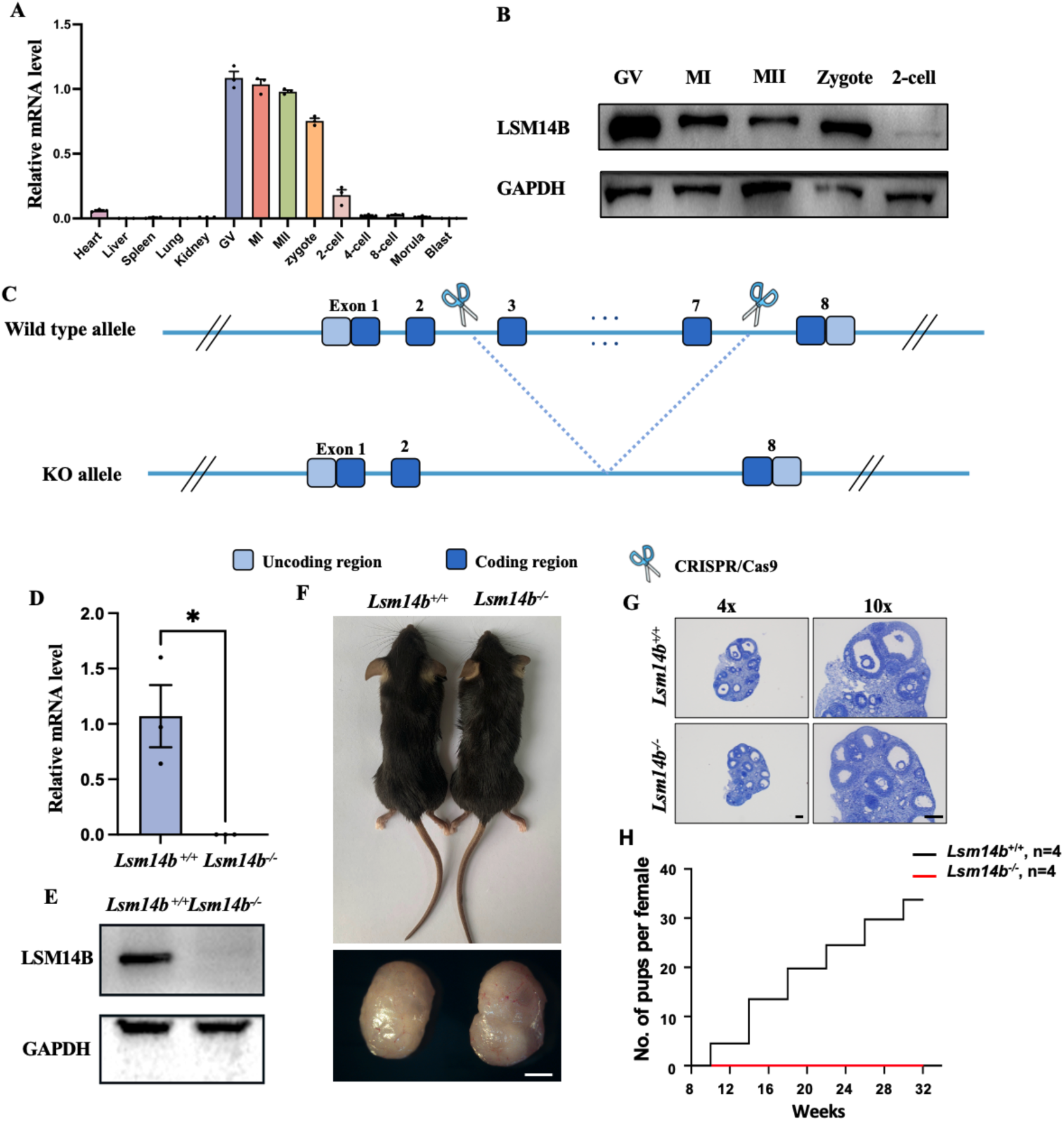
Profiling LSM14B expression in oocytes and early embryos. (A) qPCR results showing *Lsm14b* expression in mouse somatic tissues, oocytes, and early embryos. (B) Immunoblotting against LSM14B in mouse oocytes; GAPDH was used as the protein loading control. (C) The LSM14B knockout strategy in mice. (D) *Lsm14b* transcripts were detected in Wildtype but not LSM14B knockout oocytes by qPCR. Error bars, S.E.M. *P<0.05 by two-tailed Student’s t-tests.(E) Immunoblotting supporting the absence of LSM14B for the LSM14B knockout oocytes. (F) Morphology of mice and ovaries at postnatal day (PD) 56. Scale bar = 1 mm. (G) Hematoxylin staining of ovary sections from Wildtype and LSM14B knockout female mice. Scale bar = 200µm. (H) Cumulative numbers of pups per female during the indicated time windows of the fertility test. The numbers of analyzed mice are indicated (n).

We generated LSM14B knockout mouse strain using CRISPR/Cas9, based on a 695bp deletion from exons 3-7 (Figure 1C). No *Lsm14b* was detected in LSM14B knockout oocytes in qPCR (Figure 1D) and immunoblotting (Figure 1E) analyses. The body and ovary size of LSM14B knockout mice did not differ obviously from Wildtype mice (Figure 1F). LSM14B knockout female mice displayed normal ovarian histology (Figure 1G). Strikingly, fertility testing indicated that the LSM14B knockout female mice were completely infertile (Figure 1H).

### LSM14B KO oocytes display embryogenesis defects

LSM14B knockout impairs fertilization and early embryogenesis. To explore the cause of infertility LSM14B knockout female mice, we examined oocyte meiotic progression. We mated superovulated LSM14B knockout female mice with wildtype male mice to assess fertilization in detail. This produced few if any normal two-cell embryos, four-cell embryos, eight-cell embryos, or blastocysts (Figure 2A and 2D), so we conducted immunofluorescence staining on zygotes (Figure 2B). About 80% of the zygotes from LSM14B knockout mice exhibited abnormal monopronuclear (1PN) states, a significantly higher proportion than observed for zygotes from wildtype mice around 18%. In contrast, the percentage of normal bipronuclear (2PN) state zygotes from wildtype mice was 84.33%, while it was only around 22% from the LSM14B knockout mice (Figure 2C).

**Figure 2.**
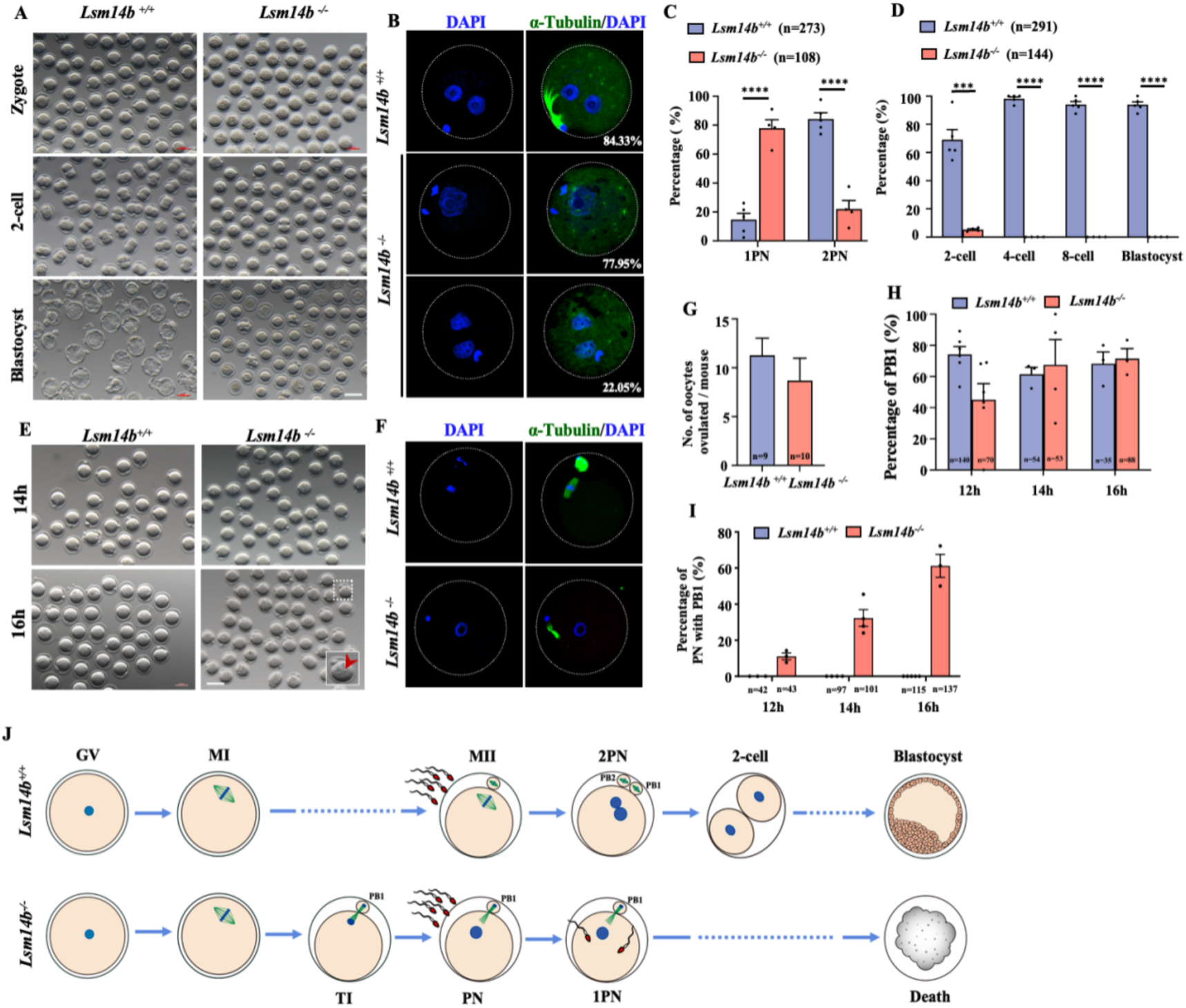
LSM14B knockout oocytes display embryogenesis defects. (A) Representative images of preimplantation embryos at different stages derived from wildtype and LSM14B knockout mice. (B) Immunofluorescence staining (as indicated) of zygotes derived from Wildtype and LSM14B knockout mice mated with wildtype males. Scale bar = 20 µm. (C) The rates one pronuclear (1PN) and 2PN zygotes collected from females mated with wildtype males. The numbers of analyzed oocytes are indicated (n). Data are presented as the mean ± SEM. \*\*\**\*p* < 0.0001.(D) Quantification of preimplantation embryos derived from wildtype and LSM14B knockout mice. The numbers of analyzed embryos are indicated (n). Data are presented as the mean ± SEM. \*\**\*p* < 0.001 and \*\*\**\*p* < 0.0001. (E) Representative images of oocytes derived from wildtype and LSM14B knockout mice in vivo at 14h and 16h after hCG injection. Scale bar = 100 µm. (F) Immunofluorescence analysis of oocytes collected from wildtype and LSM14B knockout mice at 16h after hCG injection. Scale bar = 20 µm. (G) The number of oocytes ovulated from Wildtype and LSM14B knockout mice. The numbers of analyzed mice are indicated (n). (H) The percentages of oocytes with a PB1 at 12h, 14h and 16h after ovulation induction (via hCG injection, 5IU).. The numbers of analyzed oocytes are indicated (n). (I) The percentages of oocytes with both a PB1 and a nucleus at 12h, 14h, and 16h after ovulation induction (via hCG injection, 5IU). The numbers of analyzed oocytes are indicated (n). (J) Schematic for Wildtype and LSM14B knockout oocyte meiotic maturation and early embryogenesis.

We then examined the meiosis II progression of the LSM14B knockout oocytes. We first obtained MII stage oocytes in vivo at 12, 14, and 16 hours after hCG injection. The rates of PB1 emission in the LSM14B knockout oocytes were comparable with those in the control group (Figure 2H), and the number of oocytes ovulated was also comparable (Figure 2G). To our surprise, 61.17% of the ovulated oocytes from the LSM14B knockout mice collected at 16h after hCG injection had a PB1 and an interphase-like nucleus, and 32.29% of the ovulated 14h oocytes from the LSM14B knockout mice had a PB1 and an interphase-like nucleus (Figure 2E, Figure 2I). Importantly, when we retrieved oocytes at 14h after hCG injection and cultured them for 72h in vitro, none of the oocytes from LSM14B knockout mice developed further Figure S1 A); These findings together indicate that loss of LSM14B function does not lead to parthenogenesis activation; rather, the defect in the LSM14B knockout oocytes occurs in meiosis. Immunofluorescence and confocal microscopy showed that the LSM14B knockout oocytes underwent normal PB1 emission, chromosome decondensation, and formation of interphase-like nuclei, but failed to enter meiosis II (Figure 2E, Figure2F). So although the resumption of oocyte meiosis I can occur in the absence of LSM14B, LSM14B is required for normal progression to meiosis II.

### LSM14B knockout does not disrupt parthenogenesis activation and Genome silence

Global transcriptional silencing is a highly conserved mechanism during the oocyte-to-embryo transition, which occurs in the absence of de novo transcription. In mice, transcription is globally silenced in the final stages of oocyte growth, before resuming at the two-cell embryo stage. Given that LSM14B is an RNA-binding protein (and could in theory protect mRNAs from degradation), it is reasonable to speculate mRNA stability may be reduced upon LSM14B knockout. We assessed the global transcription activities of the LSM14B knockout oocytes derived at 16h after hCG injection (using 2-cell collected from Wildtype as a positive control) using an a5’-ethynyl uridine (EU) incorporation assay. EU signals were not detected in LSM14B knockout oocytes with a PB1 and an interphase-like nucleus derived at 16h after hCG inject (Figure S1B). Thus, loss of LSM14B function did not affect transcriptional silencing in oocytes.

### The distribution and function of mitochondria is impaired in LSM14B knockout oocytes

SM14B knockout disrupts mitochondrial clustering in oocytes and impairs energy metabolism. The distribution of mitochondria are dynamic during oocyte maturation, mitochondria have the ability to migrate to areas of high energy consumption, which is crucial for correct oocyte maturation[22]. Besides, mitochondrial clustering defects have been linked to a range of devastating abnormities during oocyte maturation[14, 15]. Research has already indicated that mitochondria cluster on the periphery of spindle at metaphase I (MI) termed as ‘ring stage’ [11–13]. To investigate the impact of LSM14B deletion on mitochondrial distribution, we performed immunostaining for mitochondria with MitoTracker on live oocytes derived from wildtype and LSM14B knockout mice at different growth stages. As shown in Figure 3A, the mitochondrial distribution in GV stage oocytes with LSM14B deletion shifted from the uniform distribution around the germinal vesicle to subcortical accumulation. This phenomenon has been confirmed by comparing the intensity profiles of mitochondria signal in GV stage oocytes derived from wildtype to that in LSM14B knockout mice (Figure 3B and 3C). And the proportion of abnormal distribution of mitochondria caused by LSM14B deletion was as high as 80.86% in GV stage oocytes(Figure 3G). Consistent with previous studies of MI[20], our data shows when we depleted LSM14B by gene knockout, the mitochondrial clustering were impaired(Figure 3D). The same conclusion was drawn from the analysis of intensity profiles of mitochondria signal in MI stage oocytes derived from wildtype and LSM14B knockout mice (Figure 3E and 3F). In fact, there is 81.13% LSM14B KO MI stage oocytes showed an abnormal distribution of mitochondria, which without the formation of a mitochondrial ring (Figure 3G).

**Figure 3.**
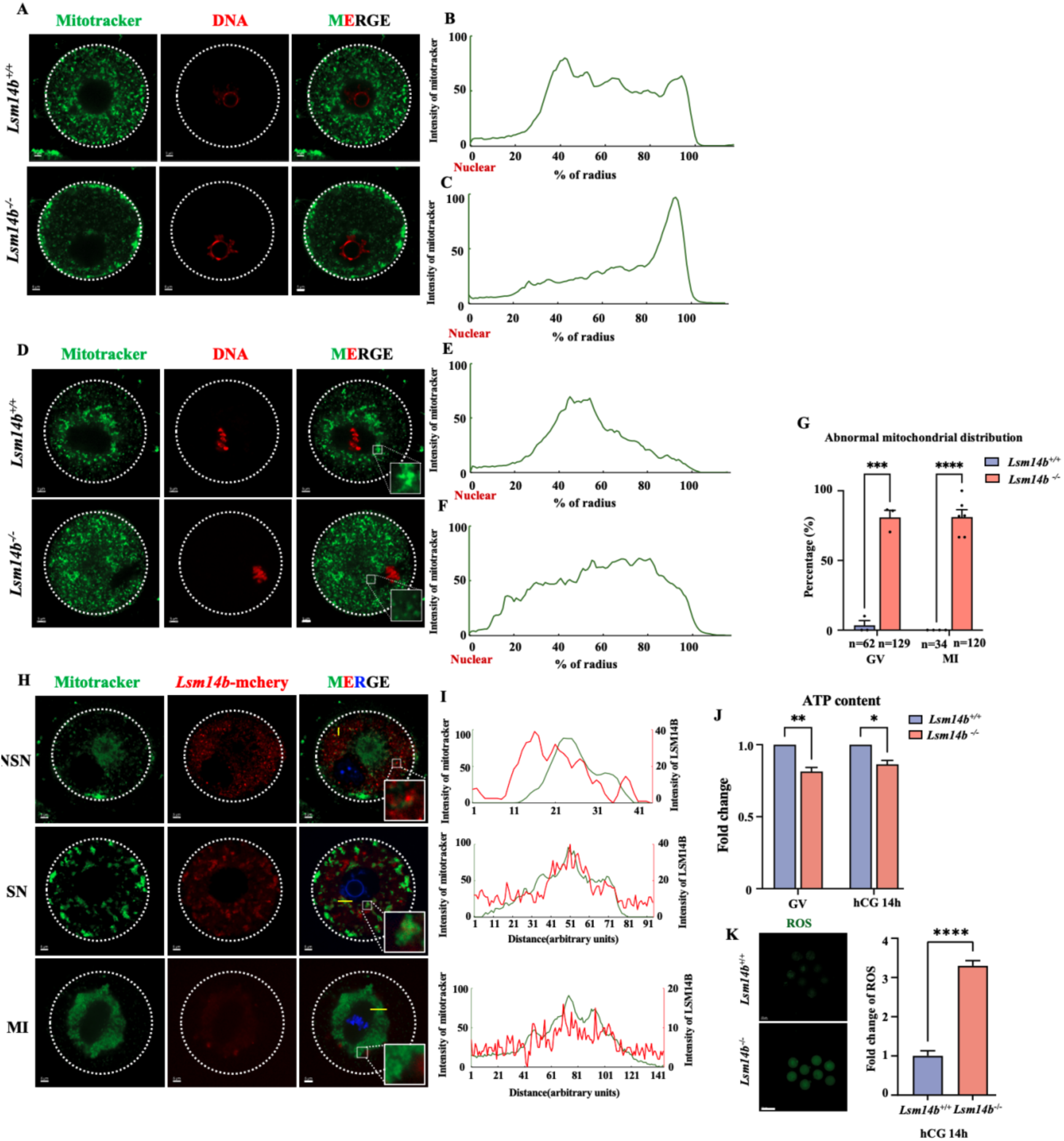
The distribution and function of mitochondria is impaired in LSM14B knockout oocytes. (A) Representative immunofluorescence images of GV stage oocytes collected from wildtype LSM14B knockout mice. The dashed line demarcates the oocyte. Scale bar = 5 µm. (B) (C) ImageJ was used to quantitatively analyze the distribution of mitochondria in panel A. (D) Representative immunofluorescence images of MI stage oocytes collected from wildtype LSM14B knockout mice. Insets are magnifications of outlined regions. The dashed line demarcates the oocyte. Scale bar = 5 µm. (E) (F) ImageJ was used to quantitatively analyze the distribution of mitochondria in panel D. (G) Rates of abnormal distribution of mitochondria in GV and MI stage oocytes. The number of analyzed embryos is indicated (n). Error bars, S.E.M. \*\**\*p* < 0.001 and \*\*\**\*p* < 0.0001 by two-tailed Student’s t-tests. (H) Representative immunofluorescence images of mouse oocytes at different growth stages. *Lsm14b*-mCherry (red), counterstained with MitoTracker (green). Insets are magnifications of outlined regions. The dashed line demarcates the oocyte. Scale bar = 5 µm. (I) Intensity profiles along the yellow lines in panel H. (J) Adenosine triphosphate (ATP) content in Wildtype and LSM14B knockout oocytes. Error bars, S.E.M. \**p* < 0.05 and \**\*p* <0.01 by two-tailed Student’s t-tests. (K) Representative images and quantification of CM-H2DCFDA fluorescence (green) in Wildtype and LSM14B knockout oocytes. Scale bar = 100 µm. Error bars, S.E.M. \*\*\**\*p* < 0.0001 by two-tailed Student’s t-tests.

To further investigate the relationship between LSM14B and mitochondria, we expressed LSM14B in the wildtype oocytes at different stages by microinjection with *Lsm14b*-mCherry mRNA and found that LSM14B co-localized with mitochondria (Figure 3H). As shown in Figure I, the co-localization of LSM14B and mitochondria reached its highest level at the MI stage, while it did not observe in NSN stage oocytes. These results suggest with the maturation of oocytes, the co-localization of LSM14B and mitochondria gradually strengthened.

Maternal mitochondria provide energy by the production of adenosine triphosphate (ATP) through the process of oxidative phosphorylation (OXPHOS) reactions for oocyte meiotic maturation and early embryonic development[11], but they also generate reactive oxygen species (ROS) which are associated with lower rates of fertilization and embryo survival[42,43]. Next, we tested the levels of ATP and ROS in the GV and MII (14h after hCG injection) stages oocytes derived from wildtype and LSM14B knockout mice. As expected, deletion of LSM14B results in a decreasing level in ATP and a increased level in ROS in both of the GV stage oocytes and MII (14h after hCG injection) stage oocytes (Figure 3J and 3K). Together, our data indicate that LSM14B has impacts on mitochondrial clustering and energy metabolism.

### LSM14B deletion altered the expression of a wide range of mitochondria-related genes

We next investigated whether the ostensibly RNA-binding protein LSM14B is associated with maternal mRNA decay. We collected fully grown GV stage oocytes from three-week old female mice at 46h after PMSG injection. For the collection of MI and MII stage oocytes, six-week old female mice were injected with 5IU human chorionic gonadotrophin (hCG) at 44h after PMSG injection, and then MI stage oocytes were collected at 8h after hCG injection, while MII stage oocytes were collected at 14h after hCG injection (Figure 4A). We performed RNA sequencing on GV and MI stage oocytes; gene expression levels were assessed as FPKM, and all samples were analyzed in triplicate. Specifically, 1042 and 892 transcripts were up- and down-regulated in LSM14B knockout GV stage oocytes; 1226 and 1453 transcripts were up- and down-regulated in LSM14B knockout MI stage oocytes (Figure 4B); qPCR verified that several RNA-binding protein and mitochondria-related transcripts decreased at both of the GV stage oocytes (Figure 4C). Further, bubble charts revealed that the transcripts decreased in LSM14B knockout GV and MI stage oocytes were primarily associated with mitochondria-related genes expression, and this phenomenon was particularly evident in LSM14B knockout MI stage oocytes (Figure 4D and 4E). The above results prompt us to consider what role LSM14B plays in the process of functional expression of mitochondria and how it affects the function of mitochondria.

**Figure 4.**
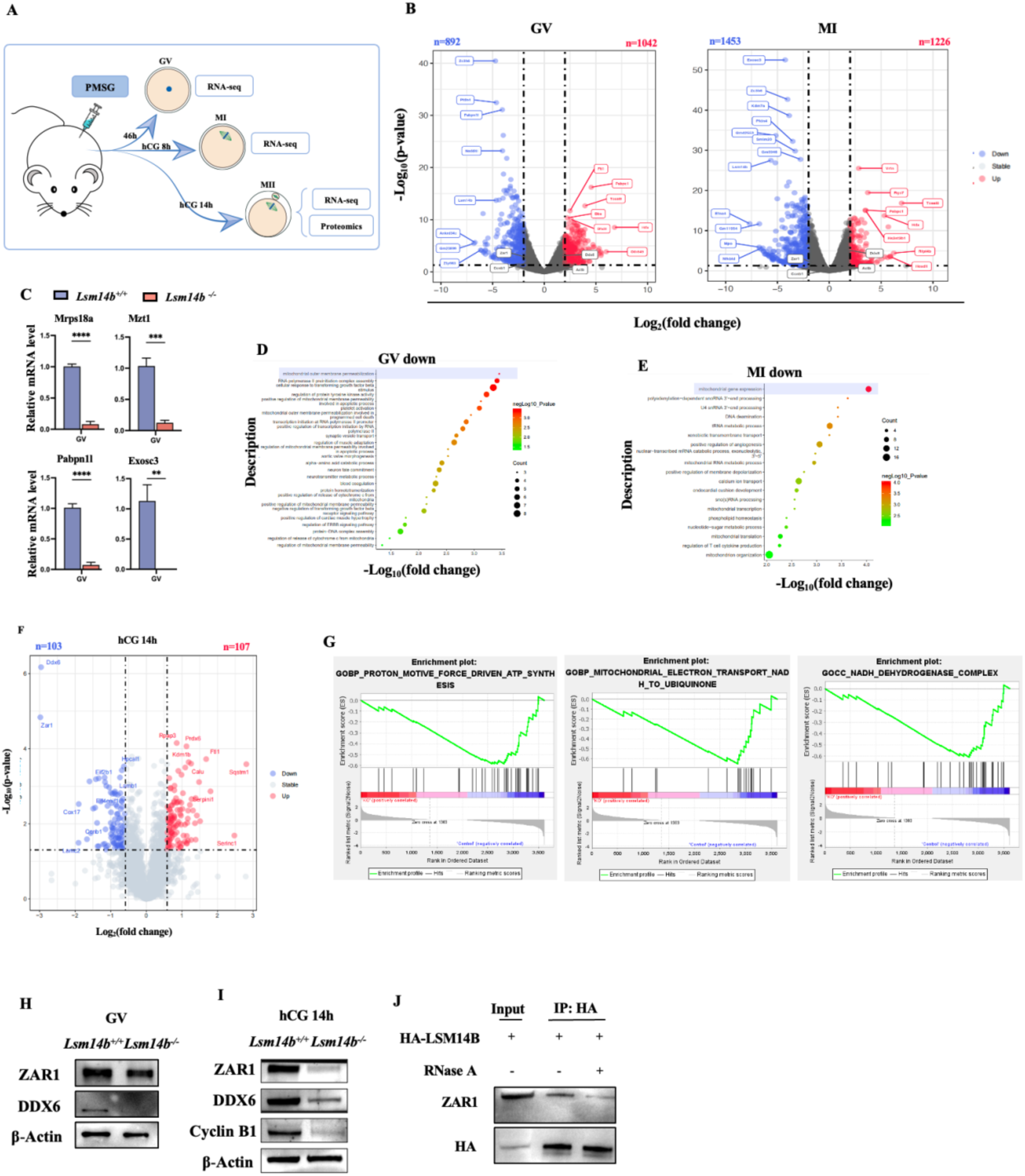
LSM14B knockout oocytes have altered expression of mitochondria-related genes. (A) Schematic showing samples for analyses of RNA-seq and proteomics. (B) Volcano plot comparing the transcripts of Wildtype and LSM14B knockout oocytes (GV, MI stage oocytes). Transcripts that increased or decreased by more than 2-fold in LSM14B knockout oocytes are highlighted in red or blue, respectively. (C) qPCR results showing expression levels of mRNAs encoding Mrps18a, Mzt1, Pabpn1l, and Exosc3 in GV stage oocytes. Error bars, S.E.M. \**\*p* < 0.01, \*\**\*p* < 0.001 and \*\*\**\*p* < 0.0001 by two-tailed Student’s t-tests. (D)(E) Bubble charts showing downregulated genes in GV and MI stage oocytes. (F) Volcano plot comparing the protein of Wildtype and LSM14B knockout oocytes collected at 14h after hCG injection. (G) The three enrichment plots from the Gene Set Enrichment Analysis (GSEA) results. (H)Immunoblotting against ZAR1 and DDX6 in GV stage oocytes derived from Wildtype and LSM14B knockout mice; β-Actin was used as the protein loading control. (I) Immunoblotting shows the expression levels of ZAR1, DDX6, and CyclinB1 proteins in MII stage oocytes (14h after hCG injection) of Wildtype and LSM14B knockout mice; β-Actin was used as the protein loading control. (J)Co-immunoprecipitation and Immunoblotting results showing the interaction between LSM14B and ZAR1. HeLa cells were transiently transfected with plasmids expressing the indicated proteins, and were harvested for co-immunoprecipitation at 48 h after plasmid transfection.

To validate this hypothesis, we collected MII stage oocytes from six-week old female mice at 14h after hCG injection (Figure 4A). A proteomics analysis detected 107 and 103 proteins that were up- and down-regulated, respectively, in LSM14B knockout MII stage oocytes (Figure 4F). Gene Set Enrichment Analysis (GSEA) revealed that pathways associated with mitochondrial electron transport, adenosine triphosphate (ATP) synthesis, and NADH dehydrogenase complex were significantly upregulated in the control group, while this phenomenon did not exesit in the LSM14B knockout group (Figure 4G). The GSEA results were consistent with the findings of the previous enrichment analysis. The known proteins DDX6, ZAR1 and Cyclin B1 exhibited particularly pronounced downregulation in the LSM14B knockout MII stage oocytes. LSM14B facilitates the initiation of MARDO assembly by binding to ZAR1 in a RNA-mediated manner. Given the proteomics results showing that the ZAR1 and DDX6 levels were significantly reduced in LSM14B knockout oocytes obtained 14h after hCG injection, we conducted immunoblotting and found that compared to MII stage oocytes, there was no significant decrease in the ZAR1 level in GV stage oocytes (Figure 4H), while the levels of both ZAR1 and DDX6 were significantly decreased in oocytes retrieved at 14h after hCG. Considering that LSM14B knockout oocytes failed to enter meiosis II, we monitored the Cyclin B1 level in oocytes obtained 14h after hCG injection. Immunoblotting demonstrated that the Cyclin B1 level was significantly decreased (Figure 4I). The complex protein Cyclin B1 regulates CDK1 activity, and is understood as the regulatory subunit. The synthesis and degradation of Cyclin B1 regulates a sequence of events during meiotic progression, including GV arrest, GVBD, the metaphase–anaphase transition of the first meiosis, and the metaphase arrest/exit of the second meiosis[44]. This knowledge, together with our observation of decreased Cyclin B1 protein levels in LSM14B knockout oocytes, implied a potential function for LSM14B in oocyte meiotic maturation. Further, we demonstrated that ZAR1 coprecipitated with LSM14B in HeLa cell, but their interaction was disrupted by RNase A treatment (Figure 4J), indicating that LSM14B and ZAR1 do not directly bind to each other but reside on the same RNA molecules. In summary, these results further supporting the likelihood that LSM14B deletion altered the expression of a wide range of mitochondria-related genes.

### Exogenous supplementation of *Lsm14b* and *Zar1* mRNA could rescue the mitochondrial distribution defects

Considering the decreased transcripts of Zar1 in LSM14B knockout oocytes, we explored whether exogenous supplement of *Lsm14b* and *Zar1* mRNA could rescue the defects of mitochondria distribution. mRNAs encoding *Zar1* and *Lsm14b* were injected into LSM14B knockout oocytes that were arrested at the GV stage using milrinone. After their release from milrinone, oocytes were stained with MitoTracker directly or cultured for 8h (Figure 5A). Consistent with the GV oocytes without exogenous mRNAs supplement, none of the distribution pattern of mitochondria was rescued in GV oocytes with microinjection of *Lsm14b* or both of *Lsm14b* and *Zar1* (Figure 5B, 5C). Conversely, mitochondria enriched at the spindle periphery in MI stage LSM14B knockout oocytes after exogenous mRNAs supplement (Figure5B, 5C). Collectively, overexpression of LSM14B and ZAR1 in oocytes significantly rescued the impaired mitochondria distribution in LSM14B knockout MI stage oocytes.

**Figure 5.**
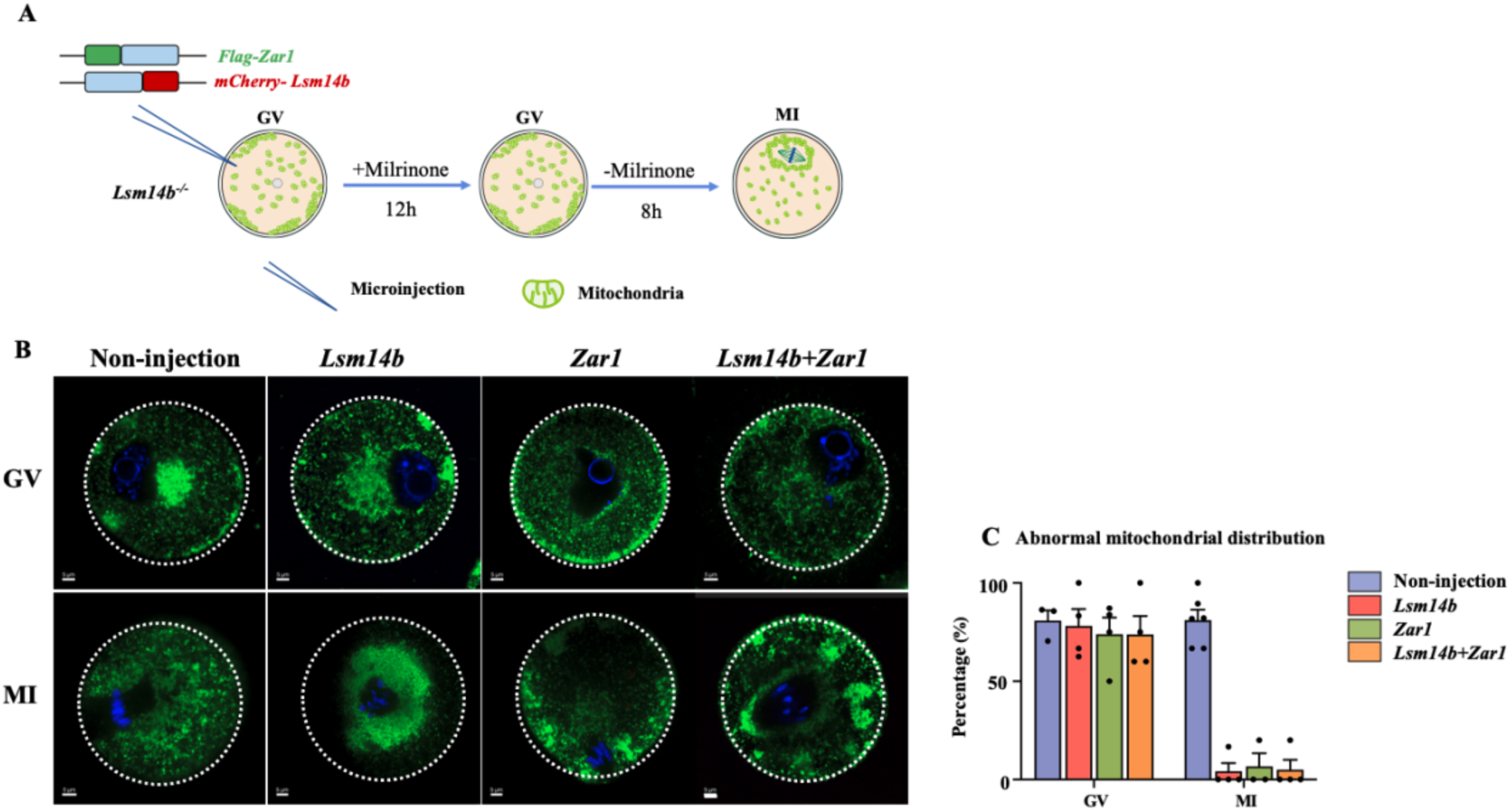
Exogenous supplementation of *Lsm14b* and *Zar1* mRNA could rescue the mitochondrial distribution defects. (A) Schematic for the rescue experimental design. (B) Representative immunofluorescence images of the mitochondria distribution in LSM14B knockoutoocytes with injection at the indicated growth stages. (C) Rates for abnormally distributed mitochondria in GV and MI stage oocytes from LSM14B knockout mice.

## Discussion

We have demonstrated that LSM14B is required for the meiosis I to meiosis II transition in mouse oocytes, and observed that female mice lacking functional LSM14B are infertile. We provide evidence that LSM14B promotes MARDO assembly in MI oocytes. We also show, through integrated analysis of transcriptomics and proteomics data (with subsequent immunoblotting validation), that LSM14B functions in regulating the translation of Cyclin B1.

Previous studies of LSM14 (of the two paralogous proteins LSM14A and LSM14B) have been conducted in *Xenopus* oocytes[16, 17]. Consistent with previous expression profile, the expression of LSM14B were specific to oocytes, indicated that LSM14B primarily functions during oocyte meiotic maturation. With LSM14B knockout mice, we have provided the first *in vivo* functional description of this gene in ensuring successive transition of meiosis I and meiosis II. In mitosis, LSM14 homologs act as RNA-binding translational repressors by binding the 3’-UTR of target mRNA to control the mitotic G2/M phase[16, 19]

A Cyclin B1-regulated clock is known to control oocyte meiotic maturation[5, 21]. Our results showed that LSM14B regulates the translational of Cyclin B1 through some post-transcriptional regulation of the *Cyclin B1* mRNA. However, in *Xenopus* oocytes, RAP55 has been be reported to represses translation when tethered to an mRNA[18]. This discrepancy may be due to the undifferentiated stage of the examined oocytes in *Xenopus*, for the translation of cyclins B1 was vary in different stage of *Xenopus* oocytes[22]. Similar to our results, Li *et al.* (2018) reported that Cyclin B1 KO oocytes entered the interphase-like phase after PB1 extrusion, without arrest at MII in mice[23]. Further experimental investigations will be required to elucidate the mechanism(s) through which LSM14B regulates the Cyclin B1 protein levels in various stages of oocyte meiotic maturation.

MARDO assembly occurs in NSN oocytes in mice, and this membraneless structure is most prominently evident in MI oocytes[24]. We found that LSM14B functions to promote assembly of MARDO in MI oocytes. Previous reports showed that ZAR1 is enriched in the cortical region of oocytes[25] and was essential for MARDO assembly and mitochondrial clustering[24]. By co-relating with ZAR1 in a RNA-mediated manner, LSM14B contributes to assembly of MARDO and the cortical distribution of ZAR1. This is reminiscent of P-bodies in *Xenopus* oocytes, and LSM14B is known to be an essential constituent for the integrity of P-bodies[26, 27]. However, with the mice oocytes grow larger P-body dispersed as the MARDO initial assemble in NSN oocytes[24, 27]. Further studies are required to research whether LSM14B is required for P-body assembly and dispersion in mice.

Mitochondria typically undergo significant changes in distribution during oocyte meiotic maturation, and localize to the periphery of the MI spindle[28–31]. Our results show that the co-localization of LSM14B with mitochondria gradually increased from the NSN to MI, and we found that supplementation with LSM14B can restore the distribution of mitochondria in LSM14B knockout mice oocytes. This result is consistent with the platform function of mitochondria for MARDO assembly[24].

## Materials and methods

### Animals

All mice were maintained in a specific pathogen free environment and all animal experiments were conducted in accordance with the guidelines and regulations of Shandong University, and the experimental protocol was approved by the Animal Care and Research Committee of Shandong University. LSM14B knockout mice were generated through CRISPR/Cas9-mediated genome engineering in a C57BL/6J background by GemPharmatech co. ltd. 695 base pairs (bp) of the *Lsm14b* gene were removed, which caused complete removal of exons 3-7. F0 founder animals were identified by PCR followed by sequence analysis and subsequently bred with wild-type mice to assess germline transmission and to generate F1 animals.

### Oocytes and embryos collection

For GV oocytes collection, mice were humanely euthanized and ovaries were dissected from three-week old female mice 46h after injection with 5 international units (IU) of pregnant mare serum gonadotrophin (PMSG). The ovaries were transferred to M2 medium (Sigma-Aldrich) and punctured with a 7-gauge needle to release cumulus–oocyte complexes, then gently remove the cumulus cells from the cumulus–oocyte complexes by a narrow-bore glass pipette.

For the collection of MI and MII oocytes, six-week old female mice were injected with 5IU human chorionic gonadotrophin (hCG) after 44h of PMSG injection and then MI-stage oocytes were collected after 8h of hCG injection, while MII-stage oocytes were collected according to specific experimental needs. Oocyte and cumulus complexes were harvested from the oviducts, digested with hyaluronidase(300IU/mL).

To obtain early embryos, superovulated female mice were mated with 12-week-old WT males for an entire night. Vaginal plugs were checked in the following morning, and mice with plugs were considered to be 0.5 days post coitum (dpc). Zygotes, two-cell, four-cell, eight-cell, morula embryos and blastocyst were collected at 1dpc, 1.5dpc, 2dpc, 2.5dpc, 3.5dpc and 4-4.5dpc respectively.

### mRNA preparation and microinjection

To prepare mRNA for microinjection, the plasmid was linearized with a restriction enzyme and then served as a template for in vitro transcription using the mMESSAGE mMACHINE T7 Ultra kit (Thermo Fisher Scientific, AM1345). Poly (A) tails [∼200 to 250 base pairs (bp)] were added to the transcribed mRNAs using the mMESSAGE mMACHINE® T7 Ultra kit. Synthesized mRNA was purified by LiCl precipitation then dissolved in nuclease-free water and quantified by NanoDrop spectrophotometer (Thermo Fisher Scientific, ND-LITE) and stored at −80°C.

All microinjections were performed using a ECLIPSE Ti2 inverted microscope (Nikon, ECLIPSE Ti2). GV oocytes were incubated in M2 medium containing 2.5μM milrinone to inhibit spontaneous germinal vesicle breakdown (GVBD) for later microinjection. Approximately 10pl of mRNAs at a concentration of 500 ng/μL were microinjected to each oocyte. After microinjection, oocytes were cultured in M16 medium containing 2.5μM milrinone for 12h at 37°C and 5% CO_2_ to ensure full translation of mRNA followed by a thorough wash and cultured in fresh M16 medium. (The plasmid of LSM14B was constructed by ourselves, while the ZAR1 was a generous gift from Dr. Qianqian Sha)

### Western blotting

For immunoblot experiments, 100 oocytes were collected and quickly washed three times with PBS, then lysed in 2×loading buffer (containing protease inhibitor). The samples were heated for 5 min at 95°C before resolved on a 12-well FuturePAGE 12% protein gel of 1.5-mm thickness (ACE Biotechnology, ET12012Gel) with MOPS-SDS running buffer (ACE Biotechnology, F00004Gel). Then proteins were transferred onto a 0.45-μm polyvinylidene fluoride membrane (Sigma Aldrich, IEVH00005) and the membranes were blocked in TBST containing 5% skimmed milk for 1 h at room temperature and then incubated with the indicated primary antibodies overnight at 4 °C. Primary antibodies against LSM14B (1/1000 dilution; Novus Biologicals, NBP2-56828), ZAR1 (1/1000 dilution; a generous gift from Dr. Hengyu Fan), DDX6 (1/1000 dilution; Proteintech, 14632-1-AP), Cyclin B1 (1/1000 dilution; CST, 4138T), GAPDH (1/1000 dilution; Proteintech, 60004-1-Ig), and β-Actin (1/1000 dilution; Proteintech, 66009-1-Ig) were used in this study. After washing three times with TBST, the membranes were incubated with the appropriate horse radish peroxidase (HRP)-conjugated secondary antibody for 1.5h at room temperature. Secondary antibodies used were HRP-conjugated goat anti-rabbit (ZSGB-BIO, ZB-2301), anti-mouse (Dako,catalog no. P0260), and anti-mouse (ZSGB-BIO, ZB-2305). All secondary antibodies were diluted 1:2000 for use. Finally, the membranes were detected by the enhanced chemiluminescence detection system (BIO-RAD, ChemiDoc MP Imaging System).

### RNA extraction and RT-PCR validation

REPLI-g WTA Single Cell Kit (QIAGEN, 150063) was used for extraction of total RNA of oocytes, and acquired ultimately cDNA according to the manufacturer’s protocols. Stored cDNA at −80℃ and diluted with RNase-free water at 1:100 for use. While RNeasy Plus Micro Kit (QIAGEN, 74034) was used for extraction of total RNA of tissues. Genomic DNA (gDNA) was eliminated by digesting with genomic DNA eraser buffer and cDNA was obtained by reverse transcription of RNA using PrimeScript RT reagent Kit with gDNA Eraser (Takara, RR047A). To measure the values of genes specifc primers, the RT-PCR was performed in triplicate. Relative mRNA levels were normalized to the level of endogenous β-actin mRNA (internal control). The relative transcription levels of the samples were compared with those of the control, followed by subsequent determination of the fold change. Finally, the relative expression levels of targeted genes were calculated using the 2^−ΔΔCT^ method. The primer sequences were as follows:

*β-actin*, 5′-AGATGTGGATCAGCAAGCAG-3′ (sense),

5′-GCGCAAGTTAGGTTTTGTCA-3′ (anti-sense);

*Exosc3,* 5′-TCGCAGCAGAAGCGGTATG-3′ (sense),

5′-ACAAAGACGCTGGCTCACTC-3′ (anti-sense);

*Pabpn1l*, 5′-CCAGTATTCAACCAGGAGTTGG-3′ (sense),

5′-GAAGCAGTCTACGGTCTCAGG-3′ (anti-sense);

*Mrps18a*, 5′-TGTTGCTCAGTCAGTTCATCC-3′ (sense),

5′-CTCCTCGATTTTTCGGTGTTCTT-3′ (anti-sense);

*Mztl*, 5′-TGGACGTTCTGCTTGAGATTTC-3′ (sense),

5′-GGCTTCTGGGTTGATTCCTTG-3′ (anti-sense).

### Histological analysis

Ovaries were collected and fixed in 4% PBS-buffered formalin overnight. After dehydration, the ovaries were embedded in paraffin and serially sectioned at 5 mm thickness and stained with hematoxylin. The slides were imaged under a microscope equipped with a camera (OLYMPUS, BX53F2).

### Immunofluorescence

Oocytes were collected and fixed in 4% PBS-buffered formalin for 30min and then permeabilized with 0.3% Triton X-100 in PBS for 20 min at room temperature. Then, oocytes were blocked in 1% BSA-supplemented PBS for 1h at room temperature followed by incubation with primary antibodies at 4°C overnight. Primary antibodies against LSM14B (1/400 dilution; Novus Biologicals, NBP2-56828), FLAG (1/400 dilution; Sigma Aldrich, F3165), α-TUBULIN-FITC (1/400 dilution; Sigma Aldrich, F2168) were used in this study. After washing three times, oocytes were incubated with appropriate secondary antibody for 1h at room temperature before stained with 4,6-diamidino-2-phenylindole (DAPI) for 10 min. Finally, oocytes were mounted on glass slides and imaged with a confocal microscope (Andor Technology Ltd, Belfast, UK).

### Visualization of the mitochondria by MitoTracker Green

MitoTracker Green (1/2000, Invitrogen, M7514) was first diluted with M16 medium to make culture droplets and placed in incubator at 37°C to balance. Oocytes were then transferred to the droplets and incubated at 37°C for 30 min. After counterstaining with Hoechst (1/800 dilution, Thermo Fisher Scientific, R37605), oocytes were moved to a glass bottom culture dish and imaged under a laser scanning confocal microscope (Andor Technology Ltd, Belfast, UK). Mitochondria distribution was reflected by MitoTracker Green staining. All photos for analysis were taken with the same intensity parameters and exposure time settings. The distribution of mitochondrial signals in a oocyte was evaluated objectively with a Clock Scan plugin for ImageJ as described previously[32].

### Determination of ATP levels

ATP testing assay kit (Beyotime, S0026) was used to measure total ATP content of oocytes. Briefly, 50 oocytes were added to 200μL lysis buffer and centrifuged at 12000×g for 5 min at 4*℃*. Supernatant was collected and mixed with testing buffer, then ATP concentration was detected on a luminescence detector (EnSpire Multimode Plate Reader). A standard curve ranging from 0.01mM to 1mM was generated and was used to calculate the total ATP contents.

### Detection of ROS generation

ROS assay kit (Beyotime, S0033S) was used to detect ROS generation in oocytes. Dichlorofluorescein diacetate (DCFH-DA) probe was diluted with M16 medium to make culture droplets and placed them in incubator at 37°C to balance for 30 min. Thereafter, oocytes were transferred to the droplets containing 10µM DCFH-DA and incubated (in the dark) for 30 min at 37°C. Then oocytes were washed 3 times and moved to a glass bottom culture dish and imaged under a laser scanning confocal microscope (Andor Technology Ltd, Belfast, UK). All photos for analysis were taken with the same intensity parameters and exposure time settings. ROS level was quantified by analyzing the fluorescence intensity of the oocytes by using the ImageJ software.

### Detection of protein synthesis

Click-iT protein synthesis assay kit (Thermo Fisher Scientific, C10428) was used to detect the total protein synthesis level of oocytes. Firstly, L-homopropargylglycine (HPG) was diluted with M16 medium to make culture droplets and placed in incubator at 37°C to balance for 30 min. Thereafter, transfer oocytes to the droplets containing 50µM HPG and incubate for 30 min at 37°C before moving the oocytes to 4% PFA for 30min and then permeabilized with 0.3% Triton X-100 in PBS for 20 min at room temperature. Next, incubate oocytes in Click-iT reaction buffer for 30 minutes at room temperature according to the instructions. After counterstaining with Hoechst 33342 (1/800 dilution, Thermo Fisher Scientific, R37605), oocytes were mounted on glass slides and imaged with a confocal microscope (Andor Technology Ltd, Belfast, UK). The HPG signal intensity represents the total protein synthesis index of the oocytes.

### EU incorporation assay

Click-iT RNA Alexa Fluor 488 Imaging Kit (Thermo Fisher Scientific, C10329) was used to detect the new RNA synthesis level of oocytes. Firstly, 5-ethynyluridine (EU) was diluted with M16 medium to make culture droplets and placed them in incubator at 37°C to balance for 30 min. Thereafter, transfer oocytes to the droplets containing 1mM EU and incubate for 1h at 37°C before fixation, permeabilization and staining with reaction buffer according to the manufacturer’s protocol. After counterstaining with Hoechst 33342, oocytes were mounted on glass slides and imaged with a confocal microscope (Andor Technology Ltd, Belfast, UK).

### RNA-seq and data analysis

Fifteen oocytes at each of the three stages (GV, MI, MII) were collected from wild-type and LSM14B knockout mice respectively, with three replicates per sample. After washing three times in phosphate-buffered saline (PBS), the samples were collected in tubes with lysis component and ribonuclease inhibitor and then transferred to −80°C freezer with liquid nitrogen. The samples were sent on dry ice to Beijing Annoroad Gene Technology Co., Ltd, which was responsible for transcriptome sequencing and data analysis. The raw data was filtered to obtain high-quality, clean data which was subsequently mapped to a reference genome using HISAT2 v2.1.0. The SAM output from this alignment was then converted into BAM format using SAMtools version 0.1.5. HTSeq v0.6.0 was used to determine the read counts mapped to each gene. Finally, expression levels of the targeted genes were evaluated using the metric of reads per kilo-base of exon transcript per million mapped reads (RPKM).

The DESeq2 package (version 1.38.3) was utilized to identify differentially expressed genes (DEGs) between the two groups (3 replicates in each group). Genes were defined as DEGs when logFC ≥ 2 and logFC ≤ −2, with a P value cutoff of 0.05. To perform Gene Ontology (GO) and pathway enrichment analyses, clusterProfiler (version 4.0.5) and Metascape were employed. For the GO enrichment analysis of differential genes, the number of genes in each GO Term was calculated according to the Gene Ontology (http://geneontology.org/), and then hypergeometric test was applied to find out that compared with the whole genomic background, Significantly enriched GO Term in differentially expressed genes. Setting calibrated P value of GO term which lower than 0.05 was different gene expression significantly enriched. Hypergeometric test was applied to the enrichment analysis of each Pathway in KEGG to find out the Pathway with significant enrichment of differentially expressed genes.

Compare the RNA-seq data from theLSM14B knockout mice with that of the wild-type mice during the different developmental stages of oocytes, and the mRNA expression levels were partially verified by real-time quantitative PCR experiments. All percentages or values from at least three biological replicates were expressed as mean ± SEM, and statistical analysis was performed with paired t-tests by GraphPad Prism 9.5.1 (GraphPad Software, San Diego, CA, USA).

### Proteomics data analysis

One hundred MII stage oocytes were collected from wild-type and LSM14B knockout mice respectively, with three replicates per sample. After washing three times in phosphate-buffered saline (PBS), the samples were collected in tubes with lysis buffer and then transferred to −80°C freezer with liquid nitrogen. The samples were sent on dry ice to Hangzhou JingJie Biotechnology Co., Ltd., which was responsible for proteomics testing and data analysis. A cutoff of the adjusted P value of 0.05 (false discovery rate–adjusted) along with a log2 fold change of 2 has been applied to determine significantly regulated proteins in WT and LSM14B knockout comparison. In this study, the Gene Ontology (GO) annotations of proteins were categorized into three groups: Biological Process, Cellular Component, and Molecular Function. To determine the significance of GO enrichment of differentially expressed proteins, Fisher’s exact test was employed with identified proteins as the background, and a P value<0.05 was regarded as significant. KEGG database was used for pathway enrichment analysis, and Fisher’s exact test was used to evaluate the significance of KEGG pathway enrichment of differentially expressed proteins with identified proteins as the background. A P value<0.05 was considered significant. Furthermore, Gene Set Enrichment Analysis (GSEA) was performed to identify specific biological pathway gene sets that were significantly associated with the expression levels between two groups of samples.

### Co-immunoprecipitation (Co-IP)

At 48h after transfection, the transfected HELA cells were quickly washed in cold PBS twice and lysed in lysis buffer (Beytime, P0013) with Protease Inhibitor Cocktail (1/25 dilution, Roche, 04693116001) at 4°C for 30 min, and then centrifuged at 12,000 × g for 15 min to harvest the supernatant. Next, extracted total proteins were incubated with target antibodies and anti-IgG Rabbit antibody overnight at 4°C. Protein A-Agarose (Roche, 11134515001) and Protein G-Agarose (Roche, 11243233001) were added to each sample and incubated for 1 hour at room temperature. Then, a portion was taken from the sample with the added target antibody was added to it and the mixture was incubated at 37℃ for 30 minutes. After being washed three times in lysis Buffer (Beytime, P0013), the coimmunoprecipitated proteins were resuspended with loading buffer and heated for 10 minutes at 95 °C for further Western blot assays.

### Statistical analysis

Data was presented as mean ± SEM of three independent experiments/samples unless otherwise specified. Two-tailed unpaired Student’s t-tests or one-way ANOVA statistical tool was applied to determine the group comparisons value when necessary. \**P* < 0.05; \*\**P* < 0.01, and \*\*\**P* < 0.001. All analysis were performed using the GraphPad Prism 9.5.1 (GraphPad Software, San Diego, CA, USA).

## Acknowledgements

We are grateful to all the participants involved in this study.

## Funding

This work was supported by grants from the National Key R&D Program of China (2021YFC2700200), Academic Promotion Programme of Shandong First Medical University (2019U001), General Research Fund from Research Grants Council of Hong Kong (14103418), Basic Science Center Program of NFSC (31988101), the Shandong Provincial Key Research and Development Program (2020ZLYS02), a fund from A-Smart Group to support CUHKSDU Joint Laboratory on Reproductive Genetics of CUHK, Major Innovation Projects in Shandong Province (2021ZDSYS16) and the Science Foundation for Distinguished Yong Scholars of Shandong (ZR2021JQ27), and Taishan Scholars Program for Young Experts of Shandong Province (tsqn202103192).

## Authors’ contributions

Z.J.C., Q.Q.S., and H.B.L. designed the study and provided their valuable contributions to the whole study. Y.L.W and S.Y. performed most of the experiments. T.T.L analyzed the data of RNA-seq and Proteomics. Y.L.C performed the LACE-seq experiments. M.Y.Z performed genotyping. All authors read and approved the final manuscript.

## Competing interests

The authors declare that they have no conflicts of interest.

---

**Supplementary Figure S1.**
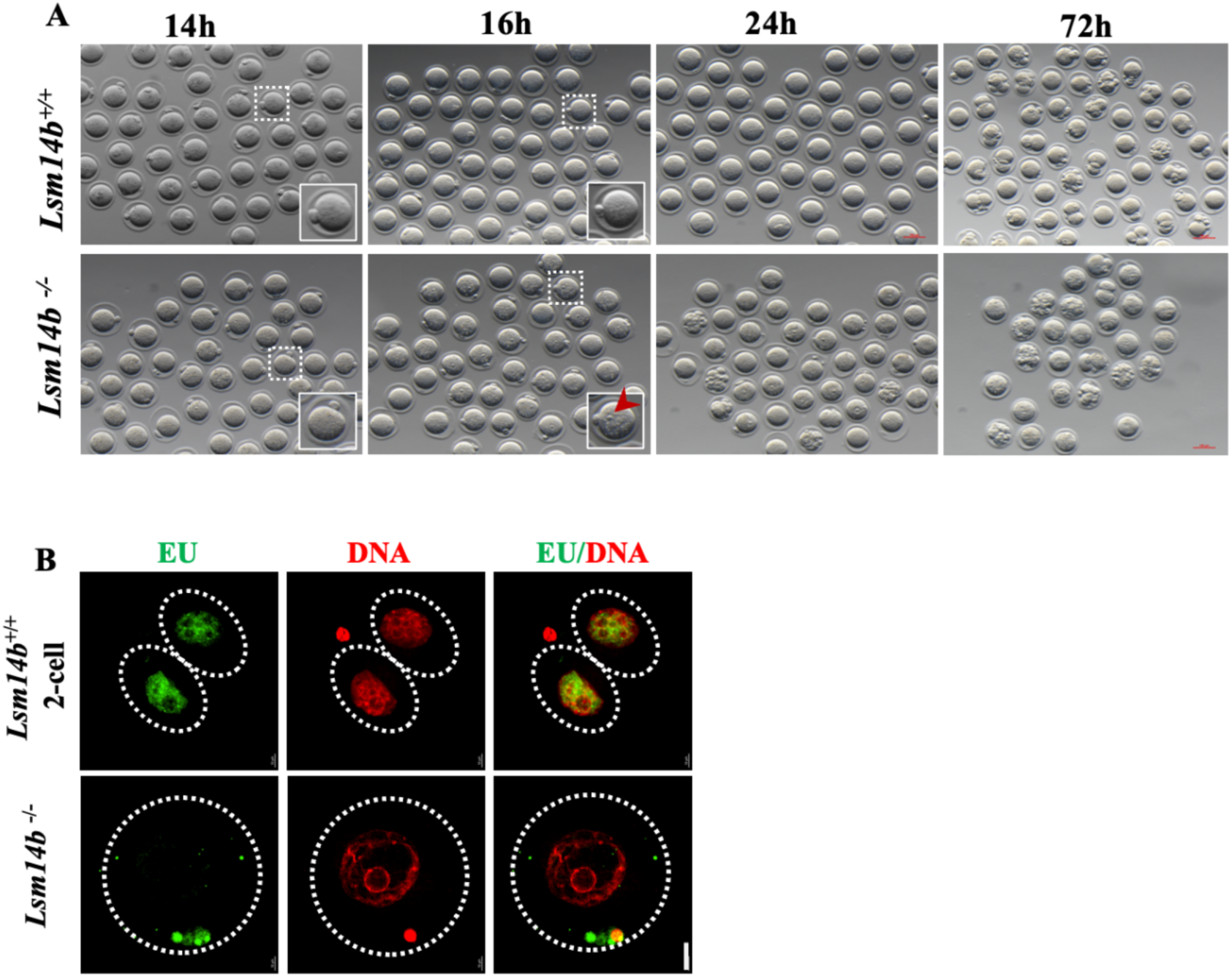
LSM14B knockout does not disrupt parthenogenesis activation and Genome silence. (A) Representative images of oocytes derived from wildtype and LSM14B knockoutmice in vitro at 14h after hCG injection. Scale bar = 100 µm. (B) Confocal images showing newly synthesized RNA by EU staining LSM14B knockout oocytes derived at 16h after hCG injection (2-cell as the positive control). Scale bar = 20 µm.

## Notes

### Competing Interest Statement

The authors have declared no competing interest.

